# Risk Factors for Chikungunya Outbreak in Kebridhar City, Somali Ethiopia, 2019. Unmatched Case-Control Study

**DOI:** 10.1101/2020.01.21.913673

**Authors:** Mikias Alayu, Tesfalem Teshome, Hiwot Amare, Solomon Kinde, Desalegn Belay, Zewdu Assefa

## Abstract

**Background:** *Chikungunya Virus* is a Ribose Nucleic Acid (RNA) virus transmitted by a mosquito bite. *Aedes Aegypti* and *Aedes Albopictus* are responsible vectors for *Chikungunya Virus* transmission. CHIKV outbreaks are characterized by rapid spread and infection rates as high as 75%. A combination of health system efforts and healthy behavior practices by the community is essential for effective control.

**Methods:** Unmatched case control study was done to identify risk factors of this outbreak. One case to two controls ratios was calculated. All cases during the study period (74 cases) and 148 controls were included in the study. Bivariate and multivariable analysis were implemented. Serum samples were tested by Real Time Polymerase Chain Reaction at Ethiopian Public Health Institute laboratory.

**Results:** A total of 74 chikungunya fever cases were reported starting from 19^th^ May 2019 to 8^th^ June 2019. Not using bed net at day time sleeping (P- value < 0.001, AOR 20.8, 95CI 6.4 – 66.7), presence of open water holding container (P- value 0.023, AOR 4, 95CI 1.2 – 13.5), presence of larvae in water holding container (P- value 0.015, AOR 4.8, 95CI 1.4 – 16.8), ill person with similar sign and symptoms in the family or neighbors (P- value <0.001, AOR 27.9, 95CI 6.5 – 120.4) and wearing not full body cover clothes (P- value 0.002, AOR 8.1, 95CI 2.2 – 30.1) were significant risk factors.

**Conclusion:** Using bed nets at day time sleeping, cover the water holding containers, wearing full body cover cloths are protective factors.

## Background

*Chikungunya Virus* (CHIKV) is an RNA virus that belongs to the *Alphavirus* genus of the *Togaviridae* family transmitted by the bite of mosquitoes *Aedes aegypti* and *Aedes albopictus*. CHIKV outbreaks are characterized by rapid spread and infection rates as high as 75%; 72%–93% of infected persons become symptomatic. The disease manifests as acute fever and potentially debilitating polyarthralgia [1].

The *Aedes Mosquitoes* breed in domestic settings such as flower vases, water-storage containers, etc. and peri-domestic areas such as construction sites, coconut shells, discarded household junk items (vehicular tyre, plastic and metal cans, etc.). Adult mosquitoes rest in cool and shady areas in domestic and peri-domestic settings and bite humans commonly during the daytime [2].

Since the first outbreak in Tanzania in 1952 Chikungunya Virus has caused outbreaks in various parts of Africa. Chikungunya Virus has been found to circulate in Eastern and Central Africa. Chikungunya fever is commonly a self-resolved disease. Whereas, patients with coexisting conditions such as cardiovascular, neurologic, and respiratory disorders or diabetes needs hospitalization. Additionally, *Chikungunya Virus* may present with bleeding when co-exist with dengue fever [3–5].

*Chikungunya Virus* is a highly contagious disease that can affect up to 70% of the total population of the outbreak affected area. The virus can easily transmit across continents and the current growing of movement of people from one country to another country as well as international trade facilitate the importation of the virus [6].

Typical presentations of *Chikungunya Virus* infection are sudden onset of fever and joint pain but sometimes it may cause severe complications including myocarditis, meningitis, encephalitis, and flaccid paralysis [7].

Frequent outbreaks of *Chikungunya Virus* infection suggest that health system efforts for vector control alone may not be sufficient for effective control. A combination of health system efforts and healthy behavior practices by the community is essential for effective control of chikungunya outbreak [8].

The prevention mechanisms for *Chikungunya Virus* are reduce human mosquito contact or eliminate vector populations. In this regard, control measures should be focused on eliminating the immature stages of the mosquitoes and their larval developmental sites [9]. This study helps to identify the potential risk factors for this chikungunya outbreak in order to implement appropriate vector control measures. This study can also be used as an information source for future planning in regard to efforts towards arboviral disease controls.

## Methods

### Study area and period

This study was conducted at Kebridahar City Administration of Somali Region. Kebridahar City is located 1006 kilometers to the East direction of Addis Ababa and 380 kilometers away from Jigjiga. The city administration has ten kebeles. The area is lowland with temperature ranged from 32°C to 40°C. According to census 2007 of Ethiopia, the city administration has a total population of 117,222. Fifty seven percent (66,817) of them were males and 12% (14,067) were children below five years of age. There were two public hospitals in Kebridahar City Administration. This outbreak investigation was conducted for one month, from 25^th^ May 2019 to 25^th^ June 2019.

### Study Design

Unmatched case control study was conducted to determine the risk factors of chikungunya fever disease outbreak and to design appropriate intervention strategies in Kebridahar City Administration of Somali Region.

### Source Population

All the residents of Kebridahar City were the source population.

### Study Population

The study population for this study were all people diseased with chikungunya fever and controls selected for the study from the source population.

### Sampling Method and Sample Size

For this study the case to control ratio was 1 to 2. All cases and two controls for each case were included. Therefore, 74 cases and 148 controls were participated in the study.

#### Selection of Cases

All individuals who full fill the case definition of chikungunya fever and who was willing to participate in the study were included in the study as a case. The cases were identified both from health facility and from the community by active case search.

#### Selection of Controls

Two controls for each case were selected from the neighbor households. One control was taken randomly by lottery method from members of the household on the right side of the case’s house and the other control was selected by the same method from the household on the left side of the case’s house.

### Case Definitions and Outbreak Threshold

#### Suspected Case

A person with acute onset of fever and severe arthralgia or arthritis not explained by other medical conditions, and who resides or has visited epidemic or endemic areas within 2 weeks before the onset of symptoms [10].

#### Confirmed Case

A suspected case with one of the following laboratory findings

- Isolation of virus from, or demonstration of specific viral antigen or nucleic acid in, tissue, blood, or other body fluid, OR
- Four-fold or greater change in virus-specific quantitative antibody titers in paired serum samples, OR
- Virus-specific IgM antibodies in serum with confirmatory neutralizing antibodies in the same or a later specimen [10].

#### Outbreak Threshold

In non-endemic area a single case of suspected Chikungunya Virus is considered as a suspected outbreak and if one case is confirmed by one of the laboratory methods it is considered as a confirmed outbreak [11].

### Operational Definitions

Breteaue index: - Number of containers which have larvae of *Aedes Mosquito* per 100 households inspected.

House index: - Percentage of households from where larvae of *Aedes Mosquito* was identified per the number of households inspected.

Container index: - Percentage of containers which have at least one larvae or pupa of *Aedes Mosquito* per the number of containers inspected.

Positive household: - A household in which at least one larvae or pupa of *Aedes Mosquito* was identified in at least one water container.

Negative household: - A household in which no larvae or pupa of *Aedes Mosquito* was identified.

Positive container: - A water holding container in which at least one larvae or pupa of *Aedes Mosquito* was found.

Negative container: - A water holding container in which larvae or pupa of *Aedes Mosquito* was not found.

Kebele: - The lowest political administration structure in Ethiopian administration system.

Epidemiologically linked: - Cases who have evidence of contact with confirmed cases.

### Data Collection

Epidemiological data were collected by face to face interview of cases and controls. Entomological data was collected by observation of water containers among selected households from four high case reporting kebeles (02, 03, 09, and 10). Larvae and pupas were collected by dipper and pipette and put them in to a well labeled cup with net covers to allow them to grow into adult mosquito. Once the adult mosquitos were grown, *Aedes Mosquito* was identified by mosquito identification key.

Regarding the human (laboratory) sample for confirmation, five serum samples were collected from suspected cases and transported to Ethiopian Public Health Institute (EPHI) Arbovirus Laboratory as per the recommended cold chain protocol. The laboratory expert from the hospital was transported the sample to EPHI as per the recommended cold chain protocol.

### Data Analysis and Presentation

After the data was cleaned and checked for completeness, entered into Epi Info Version 7.2 and exported to SPSS version 23. Descriptive analysis by person, place and time were done. Bivariate and multivariable binary logistic regressions were performed to identify risk factors for this chikungunya outbreak. The result was interpreted by using odds ratio with 95% confidence level and P-value of 0.05.

Breteaue index, house index and container index were calculated for entomological data.

To confirm the etiologic agent of the outbreak, serum samples were tested by Real Time Polymerase Chain Reaction (RT-PCR) after viral Ribose Nucleic Acid (RNA) was extracted.

### Ethical consideration

Support letter was written by Ethiopian Public Health Institute, Center for Public Health Emergency Management (PHEM) to Somali Regional Health Bureau and Kebridahar City Administration Health Office. Permission to investigate the outbreak was obtained from regional health bureau and the city health office as well as the mayor office of Kebridahar City Administration. This study is an outbreak investigation. Hence, it was not pass through ethical review process because outbreak investigation is not a planned study rather it is part of the outbreak control and prevention activity.

The interviewers had explained about the objectives, the process and the benefits of the study for each participant. Each participants were asked for their informed consent and interviewing was conducted after written consent was received from the participant. In case of interviewing children, the consent was obtained from their parents and guardians. Cases identified during data collection were sent to health facility for treatment.

## Result

### Socio demographic characteristics of respondents

A total of 222 participants, 74 cases and 148 controls, were interviewed for this study. None response rate was zero.

From the total participants, 59% (132) were female and 14% (32) were children below five years old (Table 1).

**Table 1.** Socio demographic characteristics of study participants, Kebridhar City Administration, Korahe Zone of Somali Region, Ethiopia 2019.

| Variable |  | Cases |  | Controls |  | Total |  |
| --- | --- | --- | --- | --- | --- | --- | --- |
| Sex | Female | 33 | 45% | 95 | 64% | 132 | 59% |
|  | Male | 41 | 55% | 53 | 36% | 90 | 41% |
| Age | Under 5 | 2 | 3% | 30 | 20% | 32 | 14% |
|  | 5-14 | 16 | 22% | 33 | 22% | 49 | 22% |
|  | 15-45 | 49 | 66% | 54 | 36% | 103 | 46% |
|  | 45+ | 7 | 9% | 31 | 21% | 38 | 17% |
| Educational status | No formal Education | 51 | 69% | 78 | 53% | 129 | 58% |
|  | NA | 1 | 1% | 12 | 8% | 12 | 5% |
|  | Primary school | 15 | 20% | 43 | 29% | 58 | 26% |
|  | Secondary school | 7 | 9% | 15 | 10% | 22 | 10% |
| Marital status | Married | 56 | 76% | 75 | 51% | 131 | 59% |
|  | NA | 18 | 24% | 47 | 32% | 65 | 29% |
|  | Single | 0 | 0% | 15 | 10% | 15 | 7% |
|  | Widowed | 0 | 0% | 11 | 7% | 11 | 5% |
| Occupation | Daily laborer | 11 | 15% | 0 | 0% | 11 | 5% |
|  | House wife | 24 | 32% | 68 | 46% | 92 | 41% |
|  | Merchant | 21 | 28% | 22 | 15% | 43 | 19% |
|  | NA | 0 | 0% | 12 | 8% | 12 | 5% |
|  | Student | 18 | 24% | 46 | 31% | 64 | 29% |
| Family size | Above 10 | 9 | 12% | 20 | 14% | 29 | 13% |
|  | Less than 5 | 25 | 34% | 49 | 33% | 74 | 33% |
|  | 5 to 10 | 40 | 54% | 79 | 53% | 119 | 54% |
| Total |  | 74 | 100% | 148 | 100% | 222 | 100% |

### Description of cases

A total of 74 chikungunya fever cases were reported from Kebridhar City Administration starting from 19^th^ May 2019 to 8^th^ June 2019 (Figure 1).

**Figure 1.**
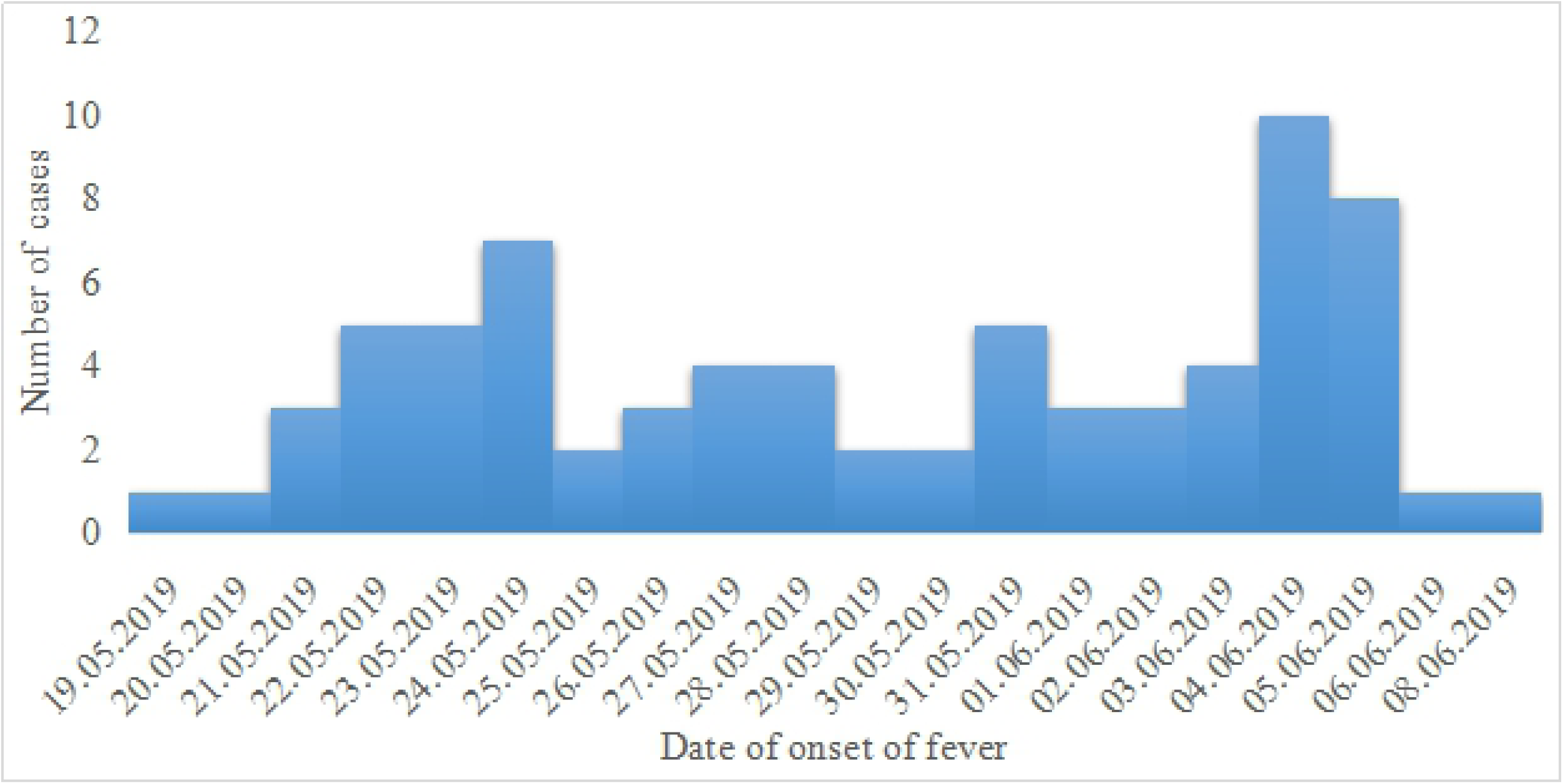
Distribution of Chikungunya fever cases by date of onset of fever in Kebridhar City Administration, Somali, Ethiopia 2019.

Of five samples sent to EPHI laboratory three were positive for Chikungunya Virus (positivity rate is 60%) and the rest of cases were epidemiologically linked. Eighty nine percent (66) were treated as an outpatient and 10.8% (12/74) cases were treated as an inpatient.

Among a total of 74 cases, 41 (55.4%) were males and 33 (44.6%) were females. Two cases were children less than five-year-old and seven cases were above 45 years old. The overall attack rate of the outbreak was 63 cases per 100,000 at risk population. The highest attack rate was among the age group of 15 – 44 (83/100,000) and the lowest attack rate was among the age group of bellow five years (22/100,000) (Table 2). The median age of cases was 25 years (IQR: 20 – 33). The case fatality rate of this outbreak was zero. The attack rates among males and females were 62 and 66 cases per 100,000 risk population respectively.

**Table 2.** Distribution of chikungunya cases by age group, Kebridhar City Administration, Somali, Ethiopia, 2019

| Age group<br>(in years) | Number of cases | Percent | At risk<br>Population | Attack Rate per<br>100,000 at risk<br>population |
| --- | --- | --- | --- | --- |
| Less than 5 | 2 | 2.70% | 9,050 | 22 |
| 5 – 14 Years | 16 | 21.60% | 35,970 | 44 |
| 15 – 44 Years | 49 | 66.20% | 59,047 | 83 |
| Above 45 Years | 7 | 9.50% | 13,155 | 53 |
| Total | 74 | 100.00% | 117,222 | 63 |

### Sign and symptoms

All cases had fever and joint pain. None of the cases had bleeding (Table 3).

**Table 3.** Distribution of chikungunya cases by sign and symptoms in Kebridhar City Administration, Somali, Ethiopia, 2019

| Sign and symptom | Yes<br>(Number) | Percent | No (Number) | Percent | Total |
| --- | --- | --- | --- | --- | --- |
| Fever | 74 | 100% | 0 | 0% | 74 |
| Joint pain | 74 | 100% | 0 | 0% | 74 |
| Headache | 53 | 72% | 21 | 28% | 74 |
| Rash | 8 | 11% | 66 | 89% | 74 |
| Nausea/vomiting | 37 | 50% | 37 | 50% | 74 |
| Bleeding | 0 | 0% | 74 | 100% | 74 |

### Entomological findings

A total of 26 household and 49 water containers were visited from four kebeles of Kebridhar City to identify the mosquito species. Among those containers in the visited households 26.5% (13/49) were positive containers and the rest 73.5% (36/49) were negative containers.

Of the visited households 38.5% (10/26) were positive households. The highest breteau index and house index were identified from Kebele Ten, whereas the highest container index was in Kebele Two of the city (Table 4).

**Table 4.** Indices of Aedes aegypti mosquito by kebele in Kebridhar City Administration, Somali, Ethiopia, 2019

| Location Investigated |  | Type of Vector Found | Aedes aegypti Stegomyia Indices |  |  |
| --- | --- | --- | --- | --- | --- |
| District | Kebeles |  | Breteau Index | House Index | Container Index |
| Kebridhar City | 02 | *Aedes aegypti* | 50% | 37.5% | 50% |
|  | 03 | *Aedes aegypti* | 28.5% | 28.5% | 22.2% |
|  | 09 | *Aedes aegypti* | 33.3% | 33.3% | 28.6% |
|  | 10 | *Aedes aegypti* | 66.6% | 44.4% | 37.5% |

### Risk factors of chikungunya fever

All participants responded that, they have bed net but their houses were not sprayed with in six months prior to this outbreak. Also, all the study participants had water holding container in their compound and they have never used mosquito repellants.

In bivariate analysis ten variables were analyzed and eight of them had a P-Value of less than 0.05. Sex, knowing the symptoms and prevention mechanisms of chikungunya, knowing that *Aedes Mosquito* bites commonly at day time, using LLINs at day time sleeping, status of water holding container, presence of larvae in water holding container, ill person with similar sign and symptoms in the family of neighbors and type of close they commonly were are significant in bivariate analysis.

In multivariable analysis eight variables significant by bivariate analysis were entered in to the model. The odds of being affected by chikungunya disease was 21 times higher among people not using LLINs at day time sleeping (P- value < 0.001, AOR 20.8, 95CI 6.4 – 66.7), four times higher among people who have open water container (P- value 0.023, AOR 4, 95CI 1.2 – 13.5), five times higher among people whose water container had mosquito larvae (P- value 0.015, AOR 4.8, 95CI 1.4 – 16.8), 28 times higher among people who have neighbors with chikungunya sign and symptoms (P- value <0.001, AOR 27.9, 95CI 6.5 – 120.4) and eight times higher among people who usually wears cloths not full body cover (P- value 0.002, AOR 8.1, 95CI 2.2 – 30.1) (Table 5).

**Table 5.** Multivariable analysis of chikungunya risk factors in Kebridhar City Administration of Somali, Ethiopia 2019

| Variables | Controls (%) | Cases (%) | P – Value | AOR | 95% CI |  |
| --- | --- | --- | --- | --- | --- | --- |
| Sex |  |  |  |  |  |  |
| Female | 95 (64.2) | 33 (44.6) |  | 1 |  |  |
| Male | 53 (35.8) | 41 (55.4) | 0.331 | 1.8 | 0.6 | 6.1 |
| Knowing symptoms and preventions of chikungunya |  |  |  |  |  |  |
| Yes | 71 (48.0) | 13 (17.6) |  | 1 |  |  |
| No | 77 (52.0) | 61 (82.4) | 0.075 | 3.3 | 0.9 | 12.0 |
| Knowing that Aedes Mosquito bites commonly at day time |  |  |  |  |  |  |
| Yes | 65 (43.9) | 21 (28.4) |  | 1 |  |  |
| No | 83 (56.1) | 53 (71.6) | 0.189 | 2.2 | 0.7 | 7.0 |
| Using LLINs at day time sleeping |  |  |  |  |  |  |
| Yes | 130 (87.8) | 12 (16.2) |  | 1 |  |  |
| No | 18 (12.2) | 62 (83.8) | <0.001 | 20.8 | 6.4 | 66.7 |
| Status of water holding container |  |  |  |  |  |  |
| Closed | 123 (83.1) | 23 (31.1) |  | 1 |  |  |
| Open | 25 (16.9) | 51 (68.9) | 0.023 | 4.0 | 1.2 | 13.5 |
| Presence of larvae in water holding container |  |  |  |  |  |  |
| Yes | 21 (14.2) | 50 (67.6) | 0.015 | 4.8 | 1.4 | 16.8 |
| No | 127 (85.8) | 24 (32.4) |  | 1 |  |  |
| Ill person with chikungunya in the house hold and neighbors |  |  |  |  |  |  |
| Yes | 15 (10.1) | 47 (63.5) | <0.001 | 27.9 | 6.5 | 120.4 |
| No | 133 (89.8) | 27 (36.5) |  | 1 |  |  |
| Kind of clothes they wear commonly since the emergence of this outbreak |  |  |  |  |  |  |
| Full body cover |  |  |  |  |  |  |
| Short | 115 (77.7) | 22 (29.7) |  | 1 |  |  |
|  | 33 (22.3) | 52 (70.3) | 0.002 | 8.1 | 2.2 | 30.1 |

## Discussion

After its reemergence in 2004, CHIKV has caused numerous epidemics around the world, including new spread to previously non endemic regions, such as the Americas, Europe, the Middle East, and Oceania [12]. The first outbreak in Ethiopia was detected in Dolo Ado District of Somali Region in 2016, since then this is the third outbreak following similar outbreak in Adar District of Afar Region in March 2019.

To confirm this chikungunya outbreak Polymerase Chain Reaction (PCR) test was done and the positivity rate was 60%. Studies in other countries also showed that, the positivity rates of CHIKV among suspected cases ranges from 12.9% to 75% [13–16].

Among the total cases 10.8% were treated as an inpatient, which is lower than the similar study in Malaysia. That study also showed that, the attack rate ranged from 0.6 to 63 per 100,000 population in different districts [17]. In our study the attack rate of the outbreak was 63 cases per 100,000 at risk population. Studies in Malaysia showed that the case fatality rate of chikungunya fever is zero [2,17]. Similarly, the case fatality rate of the chikungunya outbreak in Kebridhar City is zero.

The median age of the chikungunya fever cases in Kebridhar City was 25 years, this finding is almost similar to the median age of chikungunya fever cases (24 years) found from a cross-sectional study in Tanzania [13].

Acute phase of Chikungunya Virus infection is characterized by high grade fever and sever joint pain. Bleeding is less likely among individuals infected by Chikungunya Virus [18]. Our investigation result also supports this evidence, which founded that all cases had fever and joint pain but no case was presented with bleeding.

*Aedes Mosquito* larval index is categorized as high and low larval index based on house and bretue indices. High larval index is when the house index is ≥ 5% and/or breatue index is ≥ 20% [19]. Hence, the findings of our study shows that Bretuea index and house index of *Aedes aegypti* were ranges from 28.5% to 66.6% and from 28.5 to 44.4% respectively.

In this outbreak investigation the odds of being affected by chikungunya fever is 21 times higher among peoples who did not use bed net during day time sleep compared to those used, four times higher among peoples having open water holding container comparing to those properly close their containers, four point eight times higher among peoples whose water container had larvae of mosquitos compared to those whose water containers did not have larvae, 28 times higher among peoples living with ill persons with similar sign and symptoms compared to not living with affected peoples and eight times higher among peoples wearing shorts and T-shirts than peoples wear full body cover closes. These findings were supported by studies done in Malaysia, South India and Central Nepal. These studies shows, not using full body cover clothes, not using mosquito net or coil, having uncovered plastic water containers and staying with relatives infected with Chikungunya Virus are significant risk factors for Chikungunya Virus infection [8-9,17].

On the other hand, a retrospective study of chikungunya outbreak in India founds differences in awareness of chikungunya, cause of the disease, vector responsible, mode of transmission, biting time and elimination of breeding of mosquitoes are significant risk factors, which are not significantly associated with Chikungunya Virus infection in our study [3].

## Conclusion

This outbreak was the third chikungunya outbreak in Ethiopian by which females were more affected than males but no death was registered. Fever and joint pain were the commonest manifestation of chikungunya fever in this outbreak but no case was presented with bleeding.

This study indicated that, the larval indices of *Aedes Aegypti* mosquito was high in the city administration during the outbreak period.

Not using bed net during day time sleeping, having an open water container, presence of Aedes Mosquito larvae in water holding container, living with people having chikungunya sign and symptoms and wearing clothes which did not cover the full body were risk factors for being affected by chikungunya fever.

Therefore, there should be regular indoor and outdoor spraying of insecticidal chemicals, regular monitoring of water containers which are difficult to drain as well as to cover and apply larvicidal chemicals, awareness creation on day time bed net utilization, drainage of unusable stored water and educate the community to cover all water containers in and around the house and promote the people to wear full body cover cloths and use mosquito repellants in the time when there is an outbreak are helpful to prevent and control chikungunya outbreaks.

## Author’s contribution

MA design the study, participate in field investigation, conduct analysis and write the manuscript. TT participate in designing the study and manuscript writing. HA and DB involved in laboratory confirmation. HA and SD participate in field investigation and data analysis. All authors review the manuscript and approve the submission.

## Acknowledgement

The authors would like to acknowledge Ethiopian Public Health Institute and Somali Regional Health Bureau for facilitating the investigation and financial support during the investigation time.

